# Structural basis of connexin-36 gap junction channel inhibition

**DOI:** 10.1101/2023.12.09.570920

**Authors:** Xinyue Ding, Simone Aureli, Anand Vaithia, Pia Lavriha, Dina Schuster, Basavraj Khanppnavar, Xiaodan Li, Thorsten B. Blum, Paola Picotti, Francesco L. Gervasio, Volodymyr M. Korkhov

## Abstract

Connexin gap junction channels and hemichannels play important roles in intercellular communication and signaling. Some of connexin isoforms are associated with diseases, including hereditary neuropathies, heart disease and cancer. Although small molecule inhibitors of connexins show promise as therapeutic agents, the molecular mechanisms of connexin channel inhibition are unknown. Here, we report the cryo-EM structure of connexin-36 (Cx36) bound to an anti-malarial drug mefloquine at 2.1 Å resolution. Six drug binding sites partially occlude the pore of each connexon forming the channel. Each drug molecule in the ring makes contacts with residues in the pore-lining pocket and with the neighbouring mefloquine molecules, partially occluding the pore and modifying the pore electrostatics, ultimately reducing solute translocation through the channel. Structures of Cx36 in the presence of quinine and quinidine show a similar mode of drug binding. Molecular dynamics simulations of Cx36 bound to mefloquine show that drug binding affects the kinetics of ion passage through the pore. This previously undescribed mode of connexin channel inhibition presents an opportunity for designing subtype-specific connexin inhibitors.

**One-sentence summary:** Mechanism of connexin channel inhibition by small molecules

## Introduction

Connexins enable direct communication between cells in a tissue by forming hexameric connexons or hemichannels (HCs), which reach the cellular surface and dock with their counterparts on the neighbouring cells ^1^. The resulting assemblies, known as gap junction channels (GJCs), span two plasma membranes and couple the neighbouring cells metabolically and electrically ^2,3^. The GJCs play a crucial role in many physiological processes, such as cardiac and smooth muscle contractility (Cx43) ^4,5^, action potential propagation (Cx43) ^6,7^, neuronal synchronization (Cx36) ^8,9^, or insulin secretion (Cx36) ^10^. Different GJCs also take an active role in cell growth and differentiation, tissue development and homeostasis ^11^, and disruptions of connexin function can lead to severe pathological conditions. For example, dysfunctions in different connexin channels have been linked to deafness (Cx26) ^12^, cataracts (Cx46 and Cx50) ^13^, peripheral neuropathy (X-linked Charcot-Marie-Tooth disease, Cx32) ^14^, cardiac arrhythmias (Cx40 and Cx43) ^15,16^ and various forms of cancer ^17^.

Due to their importance in health and disease, connexins are recognized as prospective targets for therapeutic interventions. While etiology of connexin-associated diseases in many cases likely involves disrupted intercellular communication ^18-20^, blockage of connexin GJC-mediated intercellular communication or HC-mediated substrate secretion is highly desirable as a therapeutic strategy in wound healing or in cancer treatment ^21,22^. Several compounds inhibit GJCs and HCs non-specifically. For example, carbenoxolone, oleamide, anandamide, 2-APB have been shown to inhibit multiple connexin isoforms ^23-26^. It is worth noting that the binding affinity of any of these drugs for the channels has not been well established for lack of suitable tools / reagents, and much of what we know about the drug potency today is based on channel activity inhibition. Understanding the molecular mechanisms of connexin channel activity and small molecule-mediated inhibition will be essential for designing novel drugs with high specificity for different connexin isoforms, potentially offering more effective treatments to those currently available.

Cx36 is a major connexin isoform in the pancreas and in the brain ^10,27^. The physiological function of Cx36 in these tissues is to regulate neuronal activity and to mediate insulin secretion. Dysregulation of Cx36 has been associated with several human disorders. Notably, overexpression and heightened connectivity of neuronal Cx36 gap junctions has been observed in patients suffering from epilepsy, following traumatic brain injury and ischemia ^28^. The elevated gap junction coupling in these conditions contributes to subsequent neuronal death. Furthermore, in conditions such as amyotrophic lateral sclerosis (ALS), inhibiting Cx36 by pharmacological or genetic means has shown to confer neuroprotection ^29^. Therefore, development of selective inhibitors of Cx36 could be of therapeutic value. Although a cryo-EM structure of Cx36 has been solved recently, providing important insights into its molecular gating ^30^, it remains unclear how exactly drugs (or any other agents) inhibit the channel. It is also worth noting that the structural basis of drug-mediate inhibition not only in Cx36 but also in other connexins has remained unresolved to date.

Three anti-malarial drugs, mefloquine, quinine and quinidine (an enantiomer of quinine; **Figure 1a-b**), inhibit Cx36 channel with high specificity and with high apparent affinity ^31-33^. The mechanism of action of these drugs has been suggested to involve the *Plasmodium falciparum* 80S ribosome as the primary target (as in the case of mefloquine) ^34^. However, alternative research has pinpointed another intriguing target for mefloquine and quinine: *P. falciparum* purine nucleoside phosphorylase (PfPNP) ^35^. Mefloquine (commercial name: Lariam) has been shown to cause severe cardiac, neurological and psychiatric side-effects ^36-38^. Quinine (commercial name: Qualaquin) is known to cause cardiovascular side effects, blood disorders and cinchonism, a neurological disorder linked to cinchona bark intoxication (cinchona is a natural source of quinine) ^39^. Interestingly, quinidine is used clinically not only as an anti-malarial but also as an anti-arrhythmic drug (sold under commercial names Quinaglute and Quinidex), e.g. in treatment of the Brugada syndrome ^40^. Both quinine and quinidine have a variety of side effects, similar to mefloquine. The molecular mechanisms underlying the side effects of these drugs are not known and may involve multiple cellular targets, but it is likely that Cx36 inhibition and disruption of gap junction-mediated cell coupling play a role.

**Figure 1.**
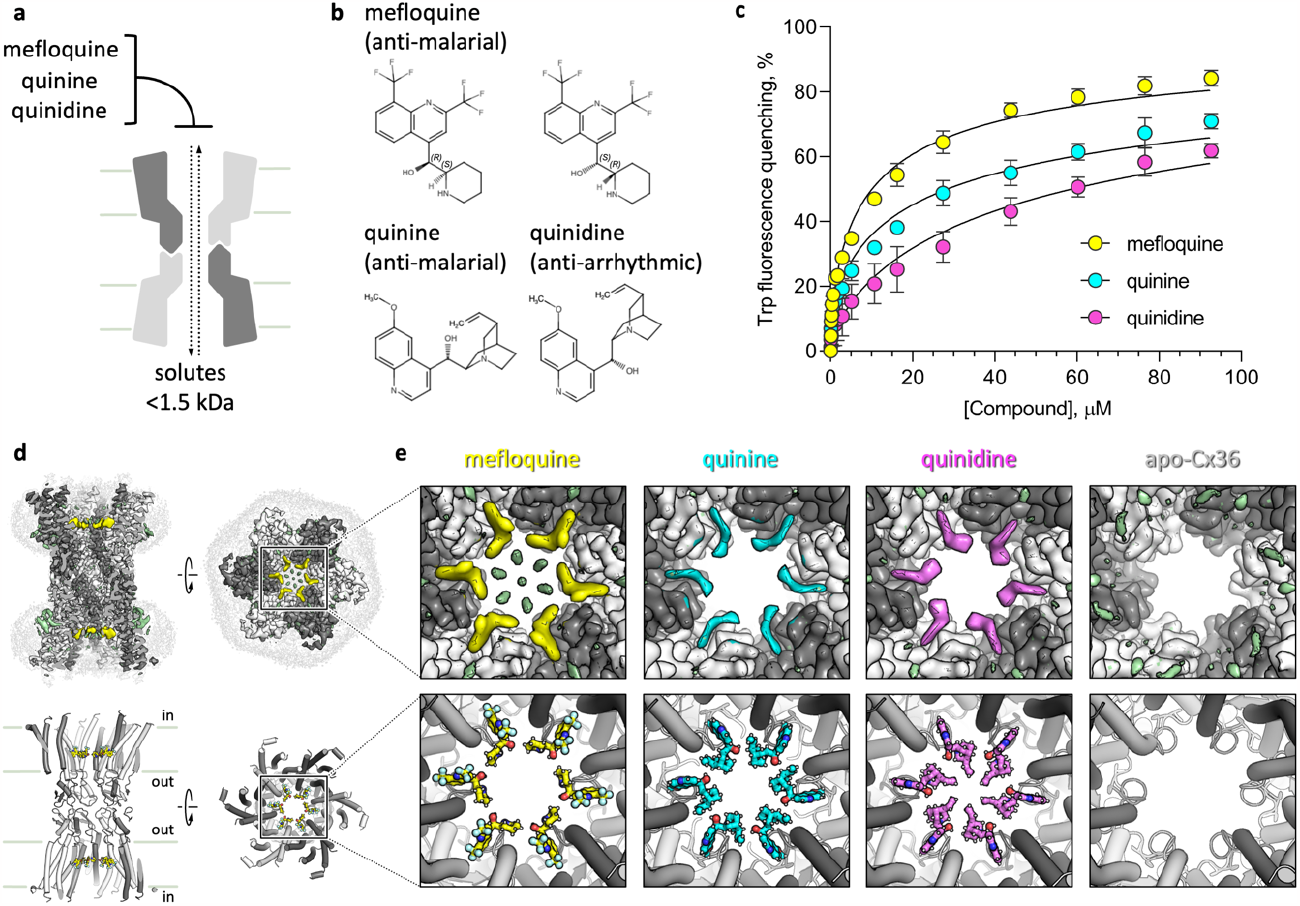
Binding of anti-malarial drugs to Cx36. **a**, A sketch illustrating the Cx36-mediated solute translocation and inhibition by selected small molecules. **b**, Chemical structures of mefloquine, quinine and quinidine. **c**, Tryptophan fluorescence quenching-based binding assays of three drugs with purified Cx36; the apparent K_d_ values are 10.4 ± 1.3 μM (mefloquine), 29 ± 5.3 (quinine), 58 ± 9 (quinidine); the values are expressed as mean ± S.D. (n = 3); the K_d_ mean values are significantly different from each other, judged by one way ANOVA. **d**, Cryo-EM map (top) and model of mefloquine-bound Cx36. Yellow density (top) corresponds to bound mefloquine; individual Cx36 monomer are coloured white and grey, to easily distinguish monomer boundaries. The light grey density corresponds to detergent micelles surrounding the membrane regions of Cx36. **e**, Zoomed-in views of maps (top) and models (bottom) of the mefloquine-, quinine- and quinidine-bound Cx36 (yellow, cyan, magenta, respectively), compared to the drug-free apo-Cx36 (right-most panels). All refined maps are contoured at 50 for comparison.

We set out to characterize the structure of Cx36 in the absence and in the presence of mefloquine, quinine and quinidine, and to determine the general principles of connexin channel inhibition by small molecules.

## Results

### Purified Cx36 binds to anti-malarial drugs *in vitro*

We expressed and purified the human Cx36 in detergent micelles (Extended Data Fig. 1a-b) and performed tryptophan fluorescence quenching-based ligand binding assays (Extended Data Fig. 1e-g, Extended Data Fig. 2). The experiments showed that all three drugs have micromolar affinity for the protein, with mefloquine binding to Cx36 with the highest apparent K_d_ value (**Figure 1c**). Experiments utilizing the nano-differential scanning fluorimetry (nano-DSF) technology showed a similar result, with an estimated K_d_ for mefloquine in the micromolar range (Extended Data Fig. 1h). These experiments confirmed that the drugs of interest interact with the purified Cx36 *in vitro* with micromolar affinity, suggesting a useful range of the drugs for our structural study.

### Structure of Cx36 in the absence and in the presence of drugs

The expressed and purified Cx36 was applied to the cryo-EM grids (in the absence or in the presence of the drugs of interest at a final concentration of 1 mM), plunge-frozen in liquid ethane and subjected to cryo-EM analysis. The acquired movies were processed as described in Materials and Methods, resulting in three reconstructions: Cx36-mefloquine at 2.14 Å resolution (Cx36-mfq; **Figure 1d-e**, Extended Data Fig. 3), Cx36-quinine at 2.73 Å resolution (Cx36-quin; **Figure 1e**, Extended Data Fig. 4), and Cx36-quinidine at 2.9 Å resolution (Cx36-quid; **Figure 1e**, Extended Data Fig. 5). The apo-Cx36 sample was prepared and analysed in a similar way, in the absence of any added compounds, resulting in a refined cryo-EM map of Cx36 at 2.49 Å resolution (apo-Cx36; **Figure 1e**, Extended Data Fig. 6, Extended Data Table 1).

The four structures show the architecture of the Cx36 GJC (**Figure 1d-e**), consistent with the recently published Cx36 in detergent and in nanodisc environment ^30^. In our case, the N-terminal gating helix is unresolved in each of the four structures (Extended Data Fig. 7b). It is possible that in the detergent environment the N-terminal helix is too dynamic to be captured by cryo-EM. Overall, the protein conformations in the three structures are nearly identical (Extended Data Fig. 7-8). The major differences were found in the pore region: the Cx36-mfq, Cx36-quin, and Cx36-quid feature an additional density, corresponding to a bound drug (**Figure 1e, Figure 2**, Extended Data Fig. 7c). Six copies of the drug are bound within the six sites in each connexon of the Cx36 GJC (**Figure 2a, c, e**).

**Figure 2.**
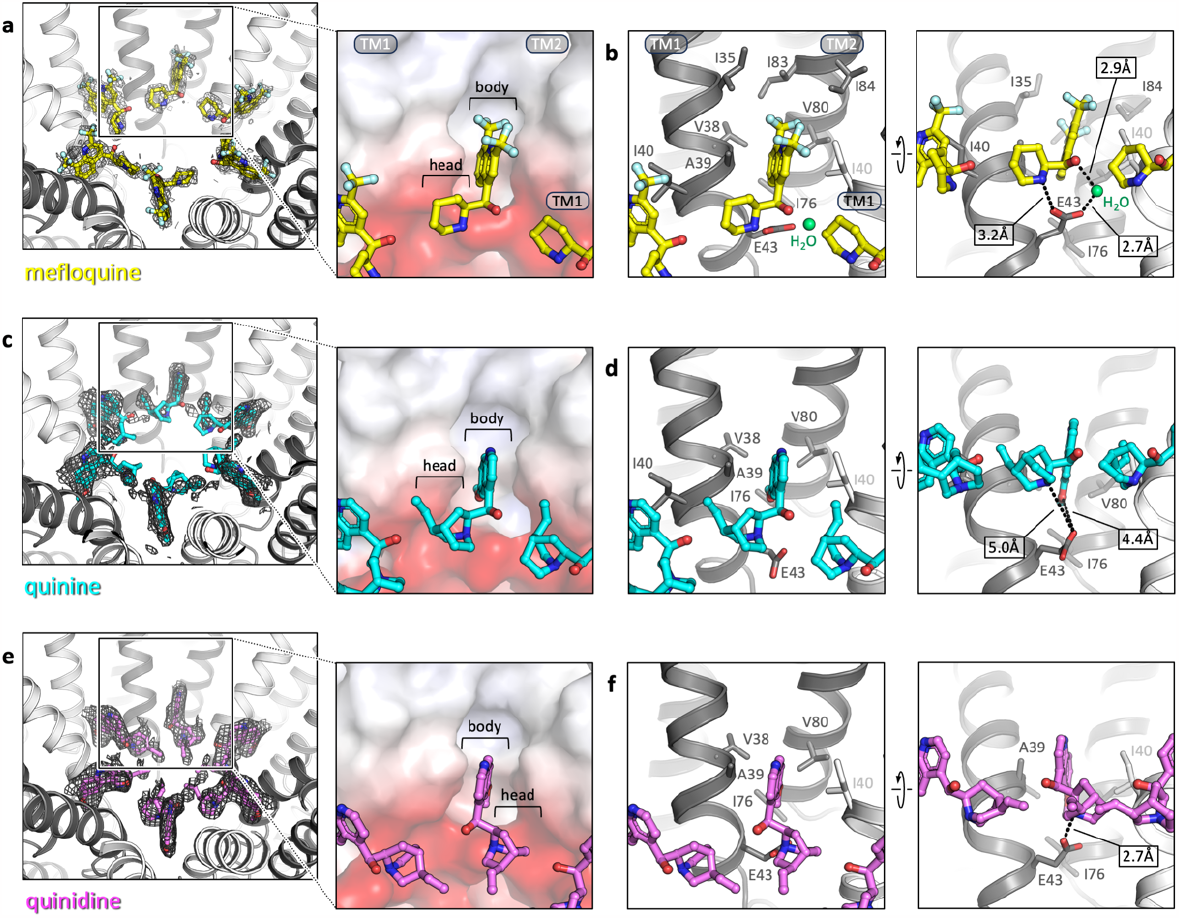
Molecular features of the inhibitor site in Cx36. **a**, Densities of six mefloquine molecules are shown as mesh (postprocessed density map, contoured at 3σ). *Inset*: the planar “body” of mefloquine is inserted into a hydrophobic groove formed by TM1 and TM2 of one Cx36 monomer and a portion of TM1 of the neighbouring Cx36 monomer; the head-group (“head”) orients towards the pore, making contacts with the polar region of the TM1 and with the neighbouring mefloquine molecule. **b**, A detailed representation of the binding site residues; a water molecule linking E43 carboxyl with the hydroxy-group of mefloquine is coloured light green. *Right*: distances (Å) between the polar atoms of mefloquine and the E43 / water are shown in white boxes. **c-d, e-f**, Same as a-b, for quinine (c-d) and quinidine (e-f).

### Details of the binding site

The structures of Cx36-mfq, Cx36-quin and Cx36-quid reveal three key elements of the drugs, relevant to connexin channel interactions: (i) a planar group that inserts into the pocket formed by residues in the transmembrane helices TM1 and TM2 (we refer to this moiety as the “body”); (ii) a hydrophobic “head-group”, which extends into the pore and makes lateral contacts with the neighbouring drug molecules; (iii) a nitrogen atom within the headgroup that contacts the conserved negatively charged residue at the periphery of the pocket (E43 in Cx36) (**Figure 2a, c, e**).

The resolution of our Cx36-mfq 3D reconstruction gives the greatest insight into the atomic details of inhibitor binding. The body of the drug wedges itself into the pocket at the interface of two Cx36 monomers, interacting with the residues I35, V38, A39 and I40 in TM1, and I76, V80, I83 and I84 in TM2 (**Figure 2b**). I40 from the neighbouring Cx36 monomer contributes to the hydrophobic pocket. The pocket is hydrophobic, and it is likely that multiple weak non-polar interactions and geometric complementarity between mefloquine and the pocket drive this interaction. The piperidinium head-group of mefloquine is asymmetrically extending toward the pore. The binding pose of mefloquine is stereoselective: although we used a racemic mixture of the drug for sample preparation, the high resolution structure captured the S,R-enantiomer of mefloquine (**Figure 2a-b**). The hydrophobic headgroup is stabilized by an interaction with the carboxyl of the residue E43 in the pore (**Figure 2b**). Moreover, the N atom of the piperidinyl group in mefloquine is within a close distance from E43 carboxyl, and the O atom of the neighbouring hydroxyl in the drug molecule is linked to E43 via an ordered water molecule (**Figure 2b**).

In the case of the quinine- and quinidine-bound Cx36, the bound drugs are somewhat less well resolved at the nominal resolution of 2.73 Å and 2.9 Å (compared to mefloquine, where the head-group ring is clearly visible in the 2.14 Å cryo-EM map). This disparity in the drug density may be attributed to the lower K_d_ of quinine/quinidine binding to Cx36 compared to mefloquine. Nevertheless, the density map quality allows us to confidently model the drugs based on the observed features (**Figure 2c, e**). In both cases the body of the drug engages in fewer hydrophobic contacts, and very distinct head-group is pointing towards the pore and making contacts with the neighbouring ligand head-group and with the E43 residue (**Figure 2d, f**). Although quinine and quinidine are stereoisomers, each appears to be readily accommodated within its binding site inside the Cx36 pore.

A combination of the hydrophobic pocket-bound ligand body, polar interactions of E43 carboxyl with the ligand head, and lateral packing of six neighbouring mefloquine molecules against each other within the Cx36 pore generates a hydrophobic ring in the translocation pathway (**Figure 3a**). Despite leaving an apparent wide aperture within the pore, the six bound mfq or quinine molecules suffice to perturb the channel function of Cx36. In the case of quinidine, the drug in the observed conformation completely closes the pore. It is worth noting that although we modelled the ligands according to the features of the experimentally determined density maps (**Figure 2c, e**), we can not exclude that alternative conformations of the ligand head-groups may exist.

**Figure 3.**
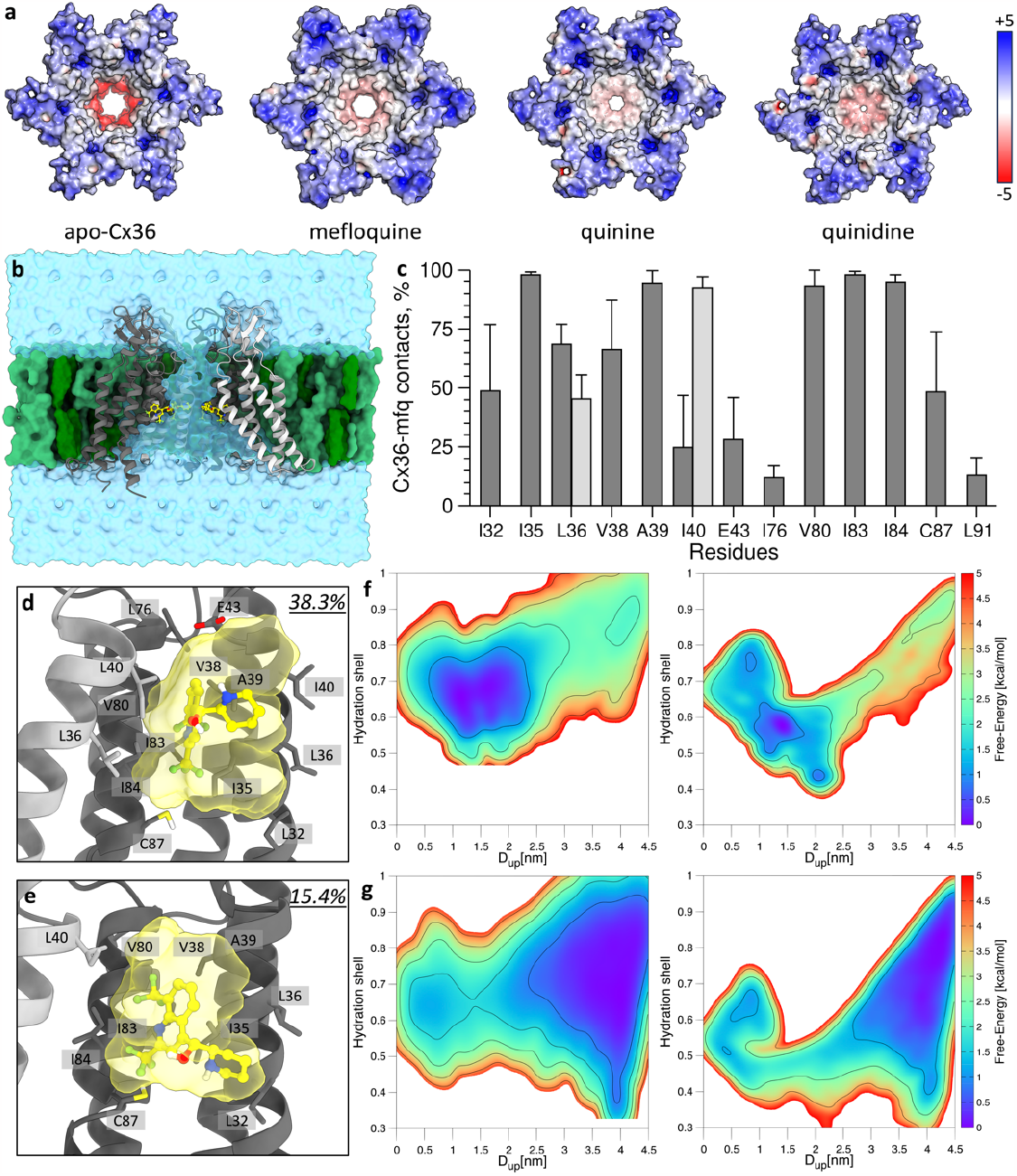
Pore electrostatics and molecular dynamics of drug-bound Cx36. **a**, The drug-free (apo-Cx36) and drug-bound structures of Cx36, in surface representation, coloured according to electrostatic potential; the scale bar corresponds to -5 / +5 kT/e. Binding of the hydrophobic drugs changes pore electrostatics (mefloquine) and/or introduces a steric barrier (quinine, quinidine). **b**, Cross-section of the membrane-embedded *6mfq-Cx36* hemichannel. To easily distinguish monomer boundaries, individual Cx36 monomer are coloured in light and dark grey. The mefloquine molecules are coloured yellow, POPC molecules - light green, and cholesterol molecules - dark green. The water content is coloured transparent cyan. **c**, Average frequencies of occurrence of the contacts between Cx36 monomers and mefloquine molecules in the *6mfq-Cx36* MD simulations. Residues belonging to different monomers are coloured in light and dark grey. The standard deviation of each point is represented through the error bar. **d-e**, Interactions established by mefloquine in cluster families *C1* (**d**) and *C2* (**e**). Their own frequency of occurrence during the *6mfq-Cx36* MD simulation is reported in the top right corners. The solvent accessible surface of mefloquine coloured transparent yellow, whereas the adjacent monomers of Cx36 are coloured dark and light grey, respectively. **f-g**, Free-energy surfaces associated with K^+^ (**f**) and Cl^-^ (**g**) permeation across the Cx36 hexamer in the *apo-Cx36* and *6mfq-Cx36* systems (left and right panels, respectively). The maps are coloured according to bars on the right side, while the isolines are placed every 1 kcal/mol.

### Small molecule binding site in other connexin isoforms

The important feature of the observed binding site is the relatively poor conservation of the residues lining the pocket, despite the structural conservation of the connexin (Extended Data Fig. 9a-b). The residues lining the drug binding site in Cx36 vary substantially in other connexins. Interestingly, in the X-ray- and cryo-EM-based 3D reconstructions of other connexin channels, this pocket is almost always filled by densities consistent with bound lipids or detergents (Extended Data Fig. 9c). While this site in distinct connexin channels interacts non-specifically with hydrophobic small molecules, our structures show that this site is used by connexin-specific drugs.

### Molecular dynamics simulations of Cx36 inhibition by mefloquine

To shed light on the dynamic behavior of the mefloquine-bound Cx36 complex and its impact on ion permeation, we performed molecular dynamics (MD) simulations. We ran 300 ns-long MD simulations of the *apo* and *holo* structures of Cx36 HC (*apo-Cx36* and *6mfq-Cx36* hereafter, **Figure 3b**). The low average Root Mean Square Deviation (RMSD) values (∼1.0 Å; Extended Data Fig. 10a) of the secondary structure Cα atoms during the simulation indicate a good conformational stability for both systems, suggesting that the presence of six ligands has a negligible impact on the overall Cx36 conformational plasticity. This was also confirmed by the per-residue Root Mean Square Fluctuations (RMSF; Extended Data Fig. 10b-c), and by a RMSD-based cluster analysis carried out on the Cx36 hexamer in the two systems (cut-off 1.5 Å). Notably, a single largely populated cluster family (containing >75% of the trajectory’s frames) was identified for both *apo-Cx36* and *6mfq-Cx36*. A superimposition of the centroids on the secondary structure’s Cαs showed a RMSD of ∼2.0 Å (see Extended Data Fig. 10d), in line with our hypothesis of negligible impact of the six ligands on the protein’s conformational plasticity.

### Dynamics of the Cx36-bound mefloquine

Having established that the structure and dynamics of Cx36 in the absence and in the presence of drug did not change dramatically, we focused on the six bound mefloquine molecules. As displayed in **Figure 3c**, mefloquine establishes predominantly hydrophobic interactions with the amino acids of two adjacent Cx36 monomers, eased by the presence of mefloquine trifluoromethyl groups (“body”). While these interactions stabilize the ligands in their own binding pockets, they still allow mefloquine to exhibit a substantial conformational freedom during the MD simulations. To quantify this behavior, we carried out a cluster analysis on each of the mefloquine molecules (cutoff 1.5 Å; see Extended Data Fig. 10e). This analysis revealed the presence of two main conformations, i.e. *C1* (**Figure 3d**) and *C2* (**Figure 3e**). While *C1* presents the mefloquine binding mode superimposable to our cryo-EM-derived structure (featuring the mefloquine “head-group” piperidinyl within a close distance from E43 carboxyl), *C2* loses the E43-mediated interaction, pointing its headgroup towards the Cx36 hexamer’s internal cavity.

### Effects of mefloquine on ion movement through the pore

To evaluate the impact of the mefloquine molecules occupying their six binding sites in Cx36 on ion translocation, we conducted a comparative analysis of the number of K^+^ and Cl^-^ ions traversing the hemichannel in both *apo-Cx36* and *6mfq-Cx36* MD simulations. As shown in Extended Data Table 2, the passage of K^+^ and Cl^-^ ions was inhibited by the drug. Over the course of our simulations, we measured 30 full K^+^ and 6 Cl^-^ transitions in the *apo-Cx36* simulation whereas only 7 K^+^ and 2 Cl^-^ were observed in the *6mfq-Cx36*. These results provide qualitative hints about the effect of the mefloquine molecules bound to their pockets in the Cx36 on ion permeation, where the ligands seem to influence the ion translocation pathway. Interestingly, the translocation pathway is not completely sterically blocked, indicating that the pore properties must be affected to cause the reduction in ion permeability.

### Free energy associated with ions permeation

To quantify the impact of mefloquine on ion permeation through Cx36, we employed a collective-variable (CV)-based free energy algorithm known as On-the-fly Probability Enhanced Sampling (OPES). This method enables the sampling of ion crossings and associated free energy profiles. Notably, OPES stands out among other CV-based enhanced sampling approaches by swiftly constructing a bias potential through on-the-fly probability estimation along selected CVs, ensuring an optimal equilibrium between exploration and convergence ^41^.

We performed two OPES simulations for 300 ns on the *apo-Cx36* system sampling the permeation of both K^+^ and Cl^-^ (*apo-Cx36_K*^*+*^ and *apo-Cx36_Cl*^*-*^, respectively). Other two 300ns-long OPES simulations were carried out on the *6mfq-Cx36* system to investigate ion translation (*6mfq-Cx36_K*^*+*^ and *6mfq-Cx36_Cl*^*-*^; detailed in the Materials and Methods and in Extended Data Fig. 11a). As shown in Extended Data Fig. 11b, the free-energy profile of K^+^ in the *apo-Cx36* and in the *6mfq-Cx36* simulations strongly differs at the level of the transition state (D_up_∼3.0 nm), where the mefloquine ligands are located. On the contrary, the free-energy of the two main basins (D_up_∼1.5 nm and D_up_∼4.0 nm) appears unaffected by mefloquine ligands, as indicated by a consistent difference of approximately 2.0 kcal/mol measured over the OPES simulation time (see Extended Data Fig. 11d). Similar scenario can also be observed for the permeation free-energy of Cl^-^, whose profile differs around the value of D_up_∼1.5-3.0 nm between the *apo-Cx36_Cl*^*-*^ and *6mfq-Cx36_Cl*^*-*^ OPES simulations (Extended Data Fig. 11c). Instead, the two basins at D_up_∼1.0 nm and D_up_∼4.0 nm are almost identical in value, and their difference along the computational time is measured to be ∼-1.7 kcal/mol (Extended Data Fig. 11e).

These results indicate that the six mefloquine molecules inhibit Cx36 by acting on the kinetics of the ion permeation instead of affecting the thermodynamics of the process. This may be due to a combined effect of (i) a modest decrease of the Cx36 pore size upon mefloquine binding, and (ii) dramatic changes in the electrostatic potential of the pore. These effects likely force the passing ions to lose their hydration shells in order to pass through the pore. To verify this suggestion, we reweighted the collected bias potential on the “Hydration shell” CV, to monitor the amount of water surrounding the ions during the crossing the Cx36 hexamer (detailed in Materials and Methods). As shown in **Figure 3f-g**, the K^+^ and Cl^-^ ion lose more than the 50% of their hydration shells while moving across the pore in the *6mfq-Cx36_K*^*+*^ *and 6mfq-Cx36_Cl*^*-*^ OPES simulations. It is worth mentioning that the Cl^-^ ions can exhibit a variety of hydration values in the intracellular side of the hexamer cavity, ranging from “fully hydrated” (∼0.9) to “poorly hydrated” (∼0.4) on a scale from 0 to 1. Such behavior, distinct from that of the K^+^ ion, seems to be due to the presence of a pronounced positive electrostatic potential on the intracellular side of Cx36 hexamer, inducing Cl^-^ ion-protein surface interactions (Extended Data Fig. 11f). Thus, while our simulations show that a complete channel blockage does not occur, consistent with the cryo-EM structure, the flux of ions is substantially reduced upon mefloquine binding.

## Discussion

Ion channel-targeted drug development is a major area of medicinal chemistry and pharmacology. Many of the drugs on the market or in phase II/III trials today target ligand- and voltage-gated ion channels, such as benzodiazepine diazepam ^42^, verapamil ^43^ and AXS-05 ^44^. The importance of ion channels for drug discovery is also exemplified by the absolute requirement to test all newly developed drugs for cross-reactivity with HERG channel to prevent cardiotoxicity ^45^. Connexin channels, including GJCs and HCs, also represent attractive drug targets and new therapies acting on Cx26 and Cx43 are currently under development ^46^. Some of the known drugs with primary action unrelated to connexins (such as mefloquine, quinine and quinidine, the topic of this study) cross-react with Cx36. This reactivity may underlie the adverse effects of these drugs. The neurological and cardiac effects of mefloquine, quinine and quinidine may well correlate with the ability of these drugs to disrupt gap junction coupling in the brain or heart, respectively. Moreover, these effects may be mediated by Cx36 or by other connexin channels that might accommodate these drugs in the corresponding drug binding sites.

The Cx36-bound drugs are sufficiently well resolved in our cryo-EM reconstructions, allowing us to get a glimpse into the atomic details of the drug-protein interaction. The mode of drug-connexin interaction relies not only on the complementarity between the drug and the protein pocket, but also on the geometry and physical properties of the drug itself. The neighbouring drug molecules within the pore interact with each other, forming a hydrophobic ring within the pore that either obstructs the pore completely or introduces the electrostatic barrier that limits solute translocation through the channel (**Figure 4**).

**Figure 4.**
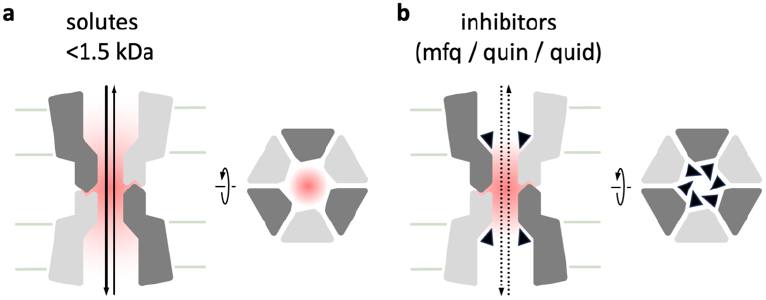
A structure-based model of drug-mediated Cx36 inhibition. **a**, Under basal conditions, Cx36 (as other connexin GJCs) is permeable to solutes with a molecular weight cut-off in the ∼1-1.5 kDa range. The electrostatic properties of the pore enable solute movement. **b**, Upon drug binding as observed in our cryo-EM 3D reconstructions (mefloquine - “mfq”, quinine - “quin”, quinidine - “quid”, shown as black triangles), steric obstruction is introduced into the path of the solutes and the electrostatic properties of the pore are altered, resulting in reduced solute flux (dotted arrows).

The mode of connexin inhibition by the small molecules described here is unique among the ion channels, and it presents an attractive new approach to tackle the challenge of drug discovery aimed at specific connexin isoforms. The following criteria would have to be satisfied to develop a connexin subtype-selective inhibitor: (i) extensive contacts and high complementary to the drug pocket (“body”); (ii) the presence of an asymmetric drug “head-group” mediating the contacts with the neighbouring drugs and with the conserve carboxyl (equivalent to E43 in Cx36). These features could build on the existing geometry and the principles of anti-malarial drug-mediated inhibition as described here, although it can be foreseen that other solutions can be found utilizing the same binding site but very different chemistry. Headgroup modification could potentially be used to fine-tune the permeability of the channels. This may be possible to accomplish due to the incomplete blockage of the channel by the drugs, evident from our structures and MD simulations. Both the strength of drug-drug interactions in the connexin pore and the permeability of the drug-bound state of the channel to small solutes or ions may be possible to tailor to the specific connexin isoforms.

## Acknowledgements

We thank Miroslav Peterek and Bilal Qureshi (ScopeM, ETH Zurich) for expert support in cryo-EM data collection. We also thank Spencer Bliven and Marc Caubet-Serrabou (PSI) for the support in high performance computing. We thank Richard Kammerer (PSI) for critical feedback on the manuscript. The work was supported by the Swiss National Science Foundation grant 184951 (VMK).

## Data and materials availability

The atomic coordinates will be deposited in the Protein Data Bank; the density maps will be deposited in the Electron Microscopy Data Bank with the following accession numbers: PDB ID 8QOJ, EMD-18540; PDB ID 8R7P, EMD-18987; PDB ID 8R7Q, EMD-18988; PDB ID 8R7R, EMD-18989. All other data are available in the main text or the supplementary materials.

## Author contributions

X.D. planned and performed the experiments, analyzed the data, wrote the manuscript, S.A. performed the simulations, analyzed the data, wrote the manuscript, A.V. performed the experiments, analysed the data, wrote the manuscript, P.L. performed the experiments, analyzed the data, D.S. performed the experiments, analyzed the data, B.K. performed the experiments, analysed the data, X.L. analysed the data, T.B.B. analysed the data, P.P. provided resources, instrumentation and methodology, F.L.G. conceptualized the study, planned and analyzed the simulations, wrote the manuscript, V.M.K. conceptualized the study, planned and performed the experiments, analyzed the data, wrote the manuscript.

## Supplementary Materials

Materials and Methods

Extended Data Figs. 1 to 11

Extended Data Tables 1 to 2

References (1-28)

