## Supplementary Information for "Structural basis of connexin-36 gap junction channel inhibition"

* Corresponding authors:

### Materials and Methods

#### Connexin-36 Expression

A synthetic gene encoding human connexin 36 (Cx36, UniprotID Q9UKL4; synthesis performed by Genewiz) with a C-terminal 3C-EYFP-twinStrep tag was cloned into the pACMV vector (REF). Freestyle 293 cells (HEK293F) cells were grown at 37°C in Dulbecco's Modified Eagle's medium (DMEM) supplemented with 10% fetal bovine serum (FBS) and then exchanged to DMEM supplemented with 2% FBS prior to transfection. The cells were transiently transfected with the expression plasmid using branched polyethyleneimine (PEI; Sigma Aldrich) at a ratio of 1:2 (w/w; DNA:PEI). After 48 hours of incubation at 37°C, cells were harvested using a cell scraper, frozen, and stored at -80°C until further use.

#### Connexin-36 Purification

Frozen cells were thawed and resuspended in buffer A (25 mM Tris-HCl, pH 8.0, 150 mM NaCl) supplemented with protease inhibitor cocktail (04693132001; Roche, Basel, Switzerland). Cells were disrupted using a Vibra-Cell sonicator, employing a 0.5-second pulse per plate, separated by a 0.5-second pause, and operated at a 35% amplitude. After sonication, the membrane fraction was clarified by ultracentrifugation (Beckman Coulter Ti45 rotor, 35,000 rpm, 50 min) and solubilized in buffer A containing 1% dodecyl-β-D-maltopyranoside (DDM) and 0.2% cholesteryl hemisuccinate (CHS) by rotating at 4°C for 1 hour. Insoluble material was removed by another round of ultracentrifugation, and the supernatant was mixed with CNBr-activated Sepharose coupled with anti-GFP nanobody and incubated at 4°C for 30 min. The resin was collected using a gravity column and washed with at least 40 column volumes of buffer B (25 mM Tris-HCl, pH 8.0, 150 mM NaCl, 0.02% glyco-diosgenin (GDN)). Cx36 was eluted overnight by addition of HRV 3C protease (0.5 mg) at 4°C and then concentrated with a 100-kDa molecular weight cutoff concentrator. Protein was injected onto a Superose 6 Increase 10/300 GL column equilibrated with buffer B for further purification. The fractions corresponding to Cx36 were collected, concentrated, and used immediately for cryo-EM grid preparation.

#### Binding assays

The binding affinity of Cx36 for ligands was assessed using intrinsic tryptophan quenching. Briefly, 5 μM of purified protein was prepared in a solution containing 25 mM Tris-HCl (pH 8.0), 150 mM NaCl, and 0.02% GDN. The protein solution was titrated with increasing amounts of ligands at sixteen titration points, ranging from 0 μM to 92 μM. Each titration point was measured using a Cary Eclipse spectrophotometer, exciting at 295 nm and recording the emission in the range of 300-500 nm. For the control titration of quinine and quinidine, drug alone in buffer was used to obtain the baseline fluorescence, which was then subtracted from the fluorescence of Cx36 in the presence of respective drug. The relative fluorescent intensity at 325 nm was plotted against the concentration of the drugs to generate the quenching profile. The apparent dissociation constant ($K_d$) values were determined by fitting the data to the specific binding with variable Hill slope model using GraphPad Prism 8.3.1.

#### Differential Scanning Fluorimetry (nano-DSF) analysis of Connexin36

A serial titration of mefloquine was mixed with an equal volume of 0.4 mg/ml Cx36 protein in a solution containing 25 mM Tris-HCl (pH 8.0), 150 mM NaCl, and 0.02% GDN. The mixture was loaded into Prometheus NT.48 Series nanoDSF Grade High Sensitivity capillaries (Nanotemper) and subjected to NanoDSF (thermal unfolding) using Prometheus Panta (Nanotemper). The temperature was continuously increased at a rate of 1°C/min from 15°C to 95°C. The raw data were analyzed with Panta Analysis software (Nanotemper) to obtain the first derivative of 350 nm/330 nm with respect to the temperature. The unfolding transition temperature ($T_m$) was determined as the maximum value of the first derivative for each titration. The obtained $T_m$ values were then plotted against mefloquine concentration and fitted with a one-site binding model using GraphPad Prism 8.3.1 software.

### Cryo-EM analysis

***Sample preparation and data collection.*** The purified Cx36 was concentrated to ~2.0 mg/mL. For the Cx36-mefloquine and Cx36-quinidine complex, a mefloquine or quinidine stock in DMSO was added to the protein to a final concentration of 1 mM. For the Cx36-quinine complex, a final concentration of 300 μM quinine in DMSO was used (at higher concentrations of the quinine the quality of the cryo-EM sample preparation deteriorated, manifesting in strongly aggregated particles judged by cryo-EM imaging as described below). The protein-drug mixture was incubated on ice for 30 min. The Quantifoil R1.2/1.3 200-mesh grids were glow-discharged for 30 s at 25 mA using a PELCO easiGlowTM glow-discharged system. A 3 μL aliquot of the protein was applied to the grid, which was then blotted and plunge-frozen in liquid ethane using a Vitrobot Mark IV (Thermo Fisher Scientific). The grids were stored in liquid nitrogen until the day of data collection. Micrographs were collected on a Titan Krios electron microscope equipped with a K3 direct electron detector and a GIF-Quantum energy filter for Cx36-mefloquine, Cx36-quinine, Cx36-quinidine and a GIF- BioContinuum energy filter for apo-Cx36 at ScopeM, ETH Zurich. The images were recorded using EPU2.0 software and dose-fractioned to 40 frames in super-resolution mode. The total dose per movie was 50, 55, 50, 55 e-/$Å^2$ for apo-Cx36, Cx36-mfq, Cx36-quin and Cx36-quid datasets, respectively.

***Cryo-EM data processing.*** The micrographs were assigned into different optics groups according to the EPU beam shift values using a script developed by Dr. Pavel Afanasyev (ETH Zurich; https://github.com/afanasyevp/cryoem_tools). The movies were corrected using MotionCor2 [1] and Gctf [2] was used for CTF estimation. For apo-Cx36, 795 particles were manually picked within Relion 4.0 [3,4]. These particles then underwent 2D classification, revealing pronounced features consistent with gap junction channels (GJCs). The most discernible 2D classes were chosen to serve as templates for the automated particle picking process across all micrographs. For the later datasets of Cx36 with drugs, the refined map of apo-Cx36 was used as template for autopicking particles. After several rounds of 2D classifications, good 2D classes were selected and extracted for 3D classification and further 3D refinement with imposed D6 symmetry. To achieve enhanced resolution, CTF refinement and Bayesian polishing were performed using Relion4.0. Additionally, the pixel size was corrected to 0.65 Å or 0.66 Å, depending on the context: apo-Cx36 or Cx36-mefloquine, Cx36-quinine and Cx36-quinidine during the postprocessing step. Local resolution maps were calculated using ResMap [5] implemented in Relion 4.0. The detailed steps of image processing are shown in Extended Data Fig. 3-6 and Extended Data Table 1.

***Model building, refinement and validation.*** The structure of apo-Cx36 was manually built in COOT [6]. The SWISS-MODEL homology model based on connexin-50 GJC (PDB ID 7JJP) of Cx36 was used as a guide for the apo-Cx36 structure. The apo-Cx36 was then used as a template for building the Cx36-mfq, Cx36-quin, Cx36-quid complexes. The cytoplasmic regions of Cx36 (M1-H18, K103-E193, A283-V321) were not built due to the poor quality of the corresponding regions in the density maps. A racemic mixture of mefloquine was added to the Cx36 protein, but only the (+)-mefloquine enantiomer (chemical ID YMZ) was found to bind to the pore of Cx36 according to the refined density map. Quinine (chemical ID QI9) from PDB 4UIL and quinidine (chemical ID QDN) from PDB 4WNU were fit into the density maps using rigid body fit. ALL the structures were refined using phenix.real_space_refine in PHENIX [7]. Model validation was performed as described previously [8]. Briefly, in order to generate FSC curve of model versus map, the coordinates of the final refined model was randomly modified 0.5 Å withing the PDB tool in Phenix. This perturbed model was subsequently subjected to refinement using one of the two available half maps. The refined model was then further iteratively refined using the other half map. The geometries of the models were validated using MolProbity [9]. All figures were prepared in PyMOL [10], Chimera [11] and ChimeraX [12].

***Electrostatic surface potential calculations***. The molecules underwent preparation for electrostatic calculations utilizing PDB2PQR [13] with the AMBER ff99 force field [14]. Determination of electrostatic surface potentials for the protein in the presence of ligands was performed using APBS Tools 2.1 [15] within PyMOL, utilizing the nonlinear Poisson-Boltzmann Equation.

***Sample preparation for LC-MS/MS analysis***

Purified protein (20 µg) was digested using a ProtiFi S-Trap™ micro spin column according to the manufacturer's protocol. The peptides were dried in a vacuum centrifuge and resuspended in 1 mL 5% acetonitrile (ACN), 0.1% formic acid (FA).

***LC-MS/MS data acquisition***

The protein samples (1 µL) were injected on a nano-flow LC system (Easy-nLC 1200, Thermo Fisher Scientific). Peptides were separated on a 40 cm x 0.75 µm (inner diameter) column packed in-house with 3 µm C18 beads at a flow-rate of 300 nL/min, a 60 min linear gradient from 3-30% II (Eluent I: 0.1% FA, Eluent II: 95% ACN, 0.1% FA) at 50°C. The samples was analyzed on an Orbitrap Exploris 480 mass spectrometer (Thermo Fisher Scientific). The samples was measured with a data-independent acquisition (DIA) method with 41 variable width DIA windows with a 1 m/z overlap. Survey MS1 spectra were recorded with a mass range between 350-1150 m/z at a resolution of 120,000 with 200% normalized AGC target or 264 ms maximum injection time. MS2 spectra covered a mass range of 150-1150 m/z at a resolution of 30,000. HCD collision energy for was set to 30% with 200% normalized AGC target or 66 ms maximum injection time.

***LC-MS/MS Data analysis***

Peptide identification and protein inference of DIA measurements was performed using Spectronaut™ software (Biognosys, version 15.5) in directDIA™ mode. Default settings were applied with minor adjustments. The minimal peptide length was set to 5 amino acids and single hits were excluded. The data were exported from Spectronaut and plots were prepared with GraphPad Prism 9.2.0. The raw file, as well as all relevant data analysis files have been deposited to the ProteomeXchange Consortium via the PRIDE [16] partner repository with the dataset identifier PXD044909.

**Molecular dynamics simulations**

***Ligand parametrization.*** Mefloquine was parametrized using Antechamber with the general Amber force field 2 (GAFF2) [17]and RESP charges fitted to ab-initio calculations with Gaussian16 following standard procedures.

***Hexamer MD simulations.*** The Cx36-mfq cryo-EM structure was used as the starting 3D structure. For each monomer, the cysteine couples *C55-C242*, *C62-C236*, and *C66-C231* have been bound with a disulfide bridge. This allowed us to obtain the following two systems: (i) *apo-Cx36*, and (ii) *6mfq-Cx36* (i.e., six Mefloquine bound inside the Connexin pore). The complexes thereby obtained were embedded into a tailored phospholipid bilayer using CHARMM-GUI [18] and solvated with TIP4P water model (salinity of 150 mM KCl). The N-terminus and C-terminus of each Connexin monomer were capped with an acetyl and a methyl-amino protecting groups, respectively. The DES-Amber force field was employed [19] in the MD engine GROMACS 2021.5 [20]. Each simulation box underwent a thermalization cycle using decreasing time-dependent restraints on heavy atoms with the following protocol: 1 ns of NVT simulation followed by 1 ns of NPT simulation for each temperature, starting from 100 K until 300 K with steps of 50 K. During the thermalization, the "V-rescale" thermostat has been employed, whereas, during the production run, we resorted to the Langevin dynamics temperature control scheme. The particle-mesh-Ewald (PME) method was used to treat the electrostatic interaction [21]. On the van der Waals interactions, a cut-off distance of 1.0 nm was applied. The pressure was fixed at a reference value equal to 1 bar thanks to the "*C-rescale*" barostat [22].

***OPES MD simulation.*** The passage of $K^+$ and $Cl^-$ ions was investigated through enhanced sampling simulation, by employing the "On-the-fly probability enhanced sampling" algorithm [23]. Two different collective variables (CVs) were used to estimate the ions' translation free-energy. To discriminate between different location of Connexin's channel, three dummy atoms have been defined along the pore, i.e., "$P_{up}$", "$P_{middle}$", and "$P_{down}$" (see Extended Data Fig 11a). For the sake of clarity, we defined $P_{up}$ as the geometric center among T51's and M52's Cα of the six Cx36's monomers, $P_{middle}$ as the geometric center among W79's Cα of the six Cx36's monomers, and $P_{down}$ as the geometric center among T20's and M21's Cα of the six Cx36's monomers.

To enhance the sampling of the ions, we selected the "Distance" CV, monitoring the distance between the $K^+$ and $Cl^-$ ions and the dummy atom $P_{up}$ (i.e., "$D_{up}$"). To avoid unphysical "jump" of the ions across the periodic boundary conditions, a harmonic restraint was placed on their distance with respect to the dummy atom $P_{middle}$. A gaussian potential was applied on $D_{up}$, with an initial value of 30 kJ/mol and a deposition rate (i.e., pace) of 500 integration steps. To run the OPES simulation, the MD engine GROMACS 2021.5 patched with PLUMED 2.7.1 was employed. Regarding the thermostat and the barostat, we used the same protocol of the unbiased MD simulations. To discriminate between different position inside the Cx36's pore, we also monitored the "*Hydration shell*" of the $K^+$ and $Cl^-$, an auxiliary CV employed to monitor the water coordination of the ions upon which we reweighted the collected bias potential. For additional information about the water coordination please refer to Refs [24,25].

***Binding interface evaluation.*** To properly assess the interactions established by the residues of Cx36 and Mefloquine, the contacts between the hexamer and the ligands have been displayed as histograms, by counting their frequency of frequency of occurrence through the PLOT NA routine of "Drug Discovery Tool" (DDT) [26]. We defined a neighboring cutoff value of 4 Å between two interacting residues.

***Cluster analysis.*** Cluster analyses on the MD trajectories were performed using GROMACS's gmx cluster routine, using the gromos algorithm. The cluster families of Cx36 in the apo-Cx36 and 6mfq-Cx36 MD simulations were obtained by aligning the trajectory on the Cα atoms of Cx36's secondary structure elements and computing the RMSD among the same sample of atoms. The cluster families on Mefloquine in the 6mfq-Cx36 MD simulation were obtained by aligning the trajectory on the Cα atoms of Cx36's secondary structure elements and computing the RMSD on Mefloquine's heavy atoms. The RMSD threshold value of 1.5 Å was chosen considering the number of cluster families generated and the similarity of protein conformations within a cluster family.

### Supplementary Figures

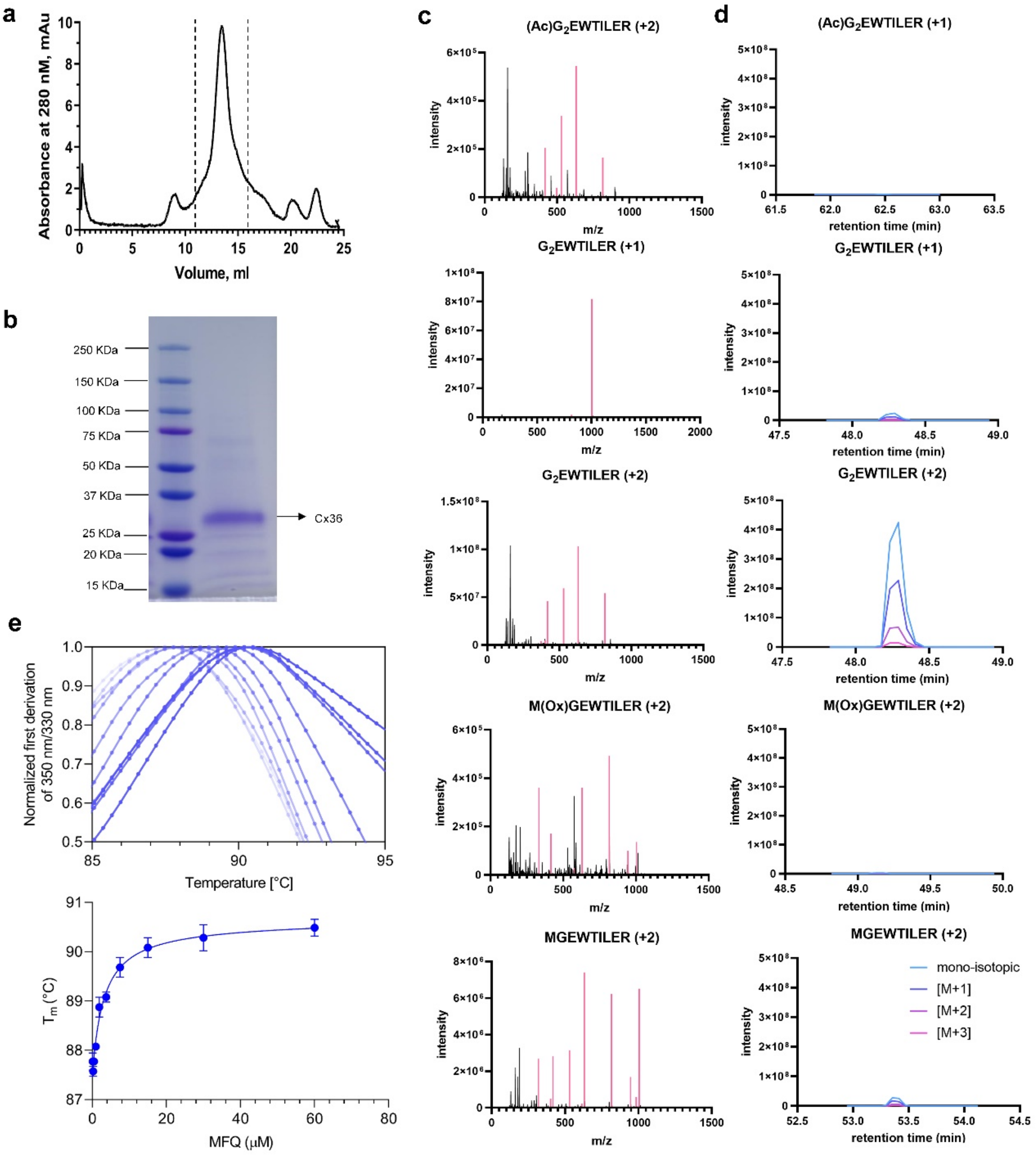

**Extended Data Fig. 1. Expression, purification and mass spectrometric characterization of Cx36.** **a,** Size exclusion chromatogram (SEC) of Cx36 purification. **b,** SDS-PAGE of Cx36 purification. **c-d,** Mass spectrometric characterization of Cx36. Fragment spectra for each of the identified peptides **(c)**. Extracted-ion chromatograms (XIC) for fragment and parent ions detected in MS2 and MS1 spectra **(d)**. **e,** Thermal unfolding (nano-DSF) graphs of Cx36 upon treatment with mefloquine. Thermal unfolding data is represented as normalization of the first derivative of the ratio of fluorescence intensity at 330 nm and 350 nm. The unfolding transition temperature (Tm), the temperature at which 50% of the protein is unfolded, is plotted against the concentration of mefloquine and fitted with a one-site binding model using GraphPad Prism software. Data is represented as mean ± SEM (n=3).

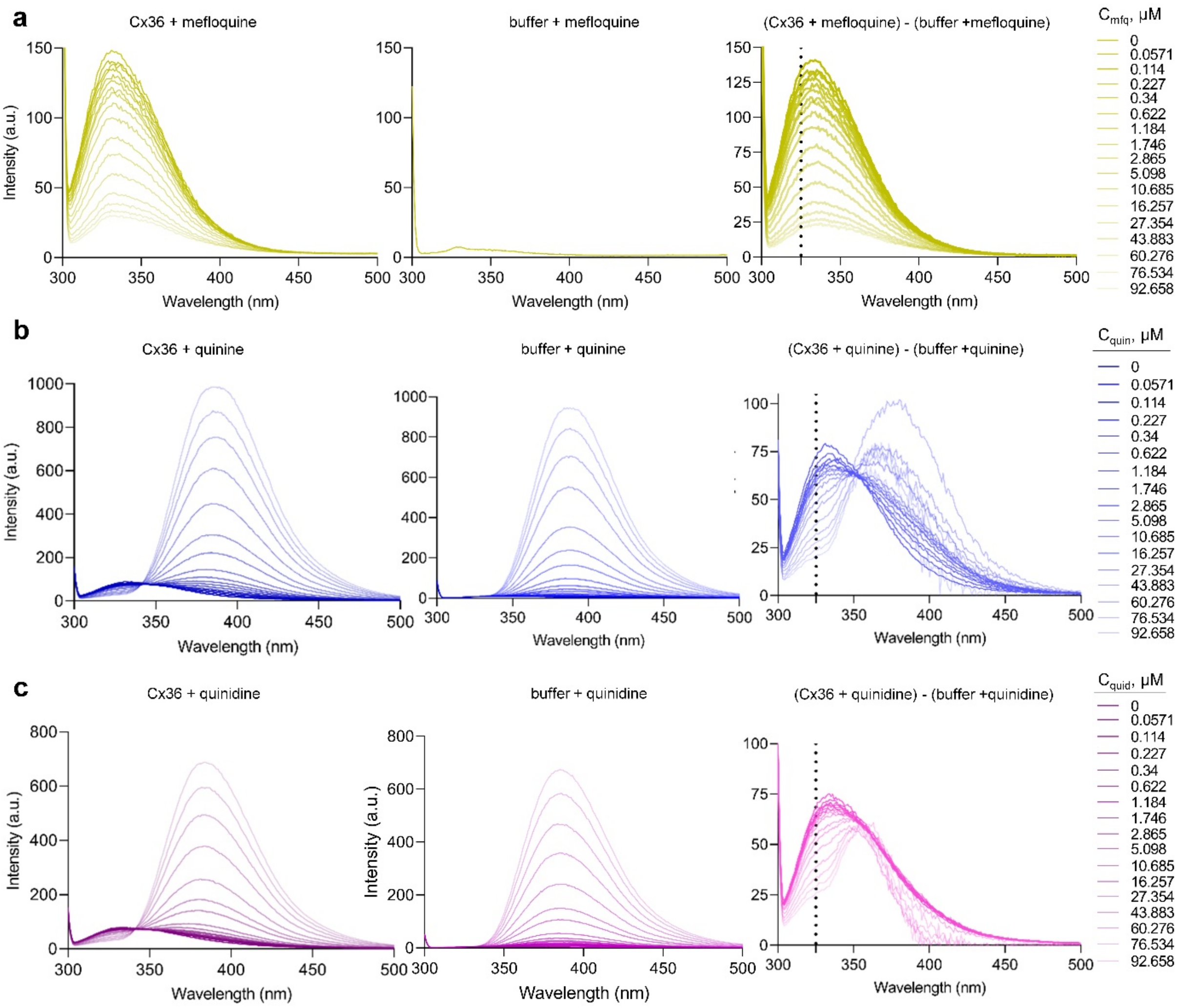

**Extended Data Fig. 2. Tryptophan fluorescence quenching of Cx36 in response to treatment with different drugs: mefloquine (a), quinine (b), quinidine (c).** The left panels display the fluorescence of Cx36 in the presence of varying concentrations of each drug, while the middle panels show the fluorescence in buffer containing different concentrations of the respective drug. In the case of mefloquine, the middle panel exclusively displays the fluorescence in the presence of 100 µM mefloquine within the buffer (a). The right panels illustrate the subtraction of these two fluorescence signals. Dot lines indicate the wavelength used for plotting the binding curves. Data is represented as mean (n=3).

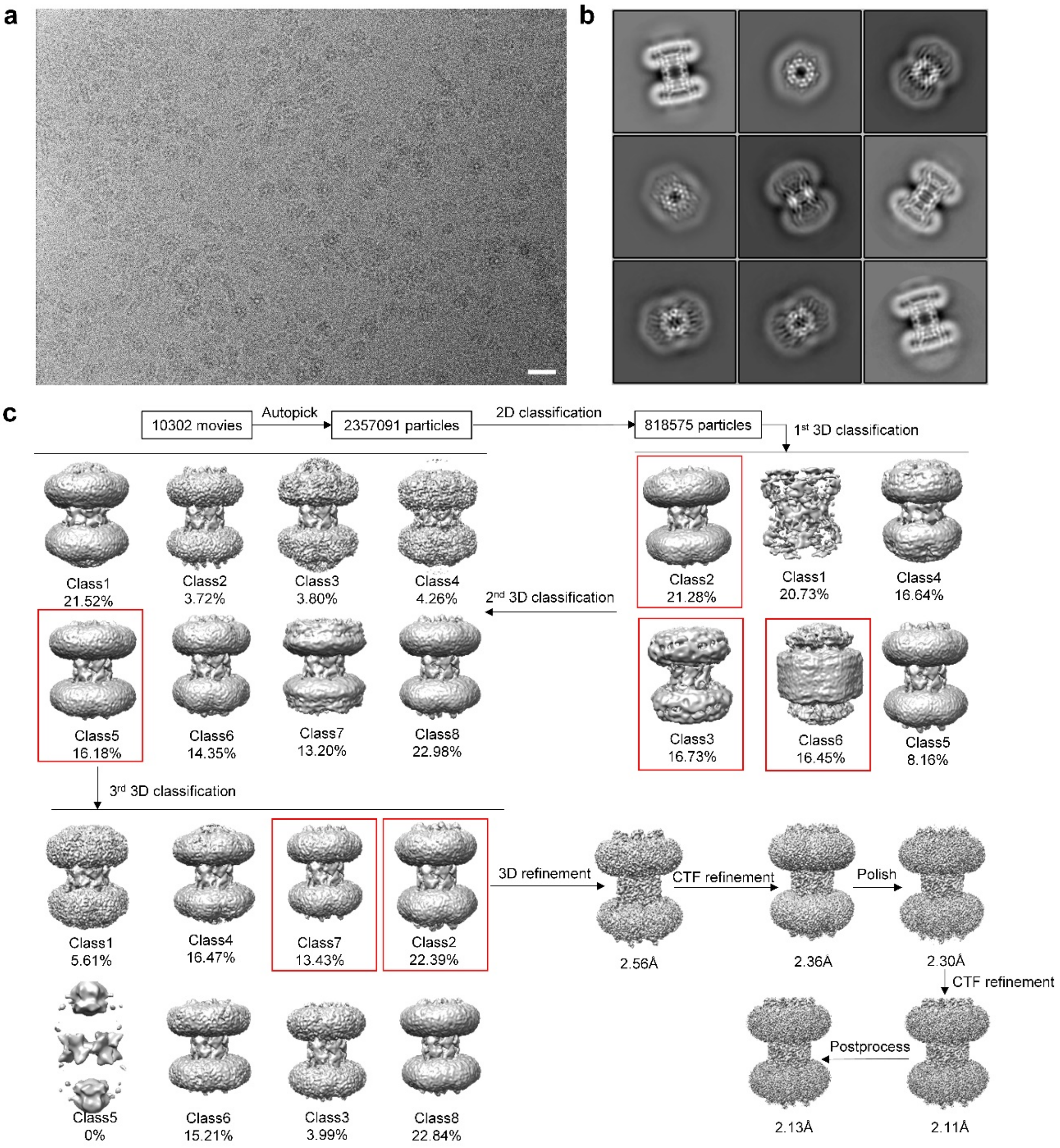

**Extended Data Fig. 3. Cryo-EM data processing pipeline of Cx36-mfq in GDN. a,** A representative micrograph of Cx36-mfq sample; the scale bar corresponds to 20 nm. **b,** representative 2D classes. **c,** Cryo-EM data process scheme for 3D reconstitution of Cx36-mfq in GDN.

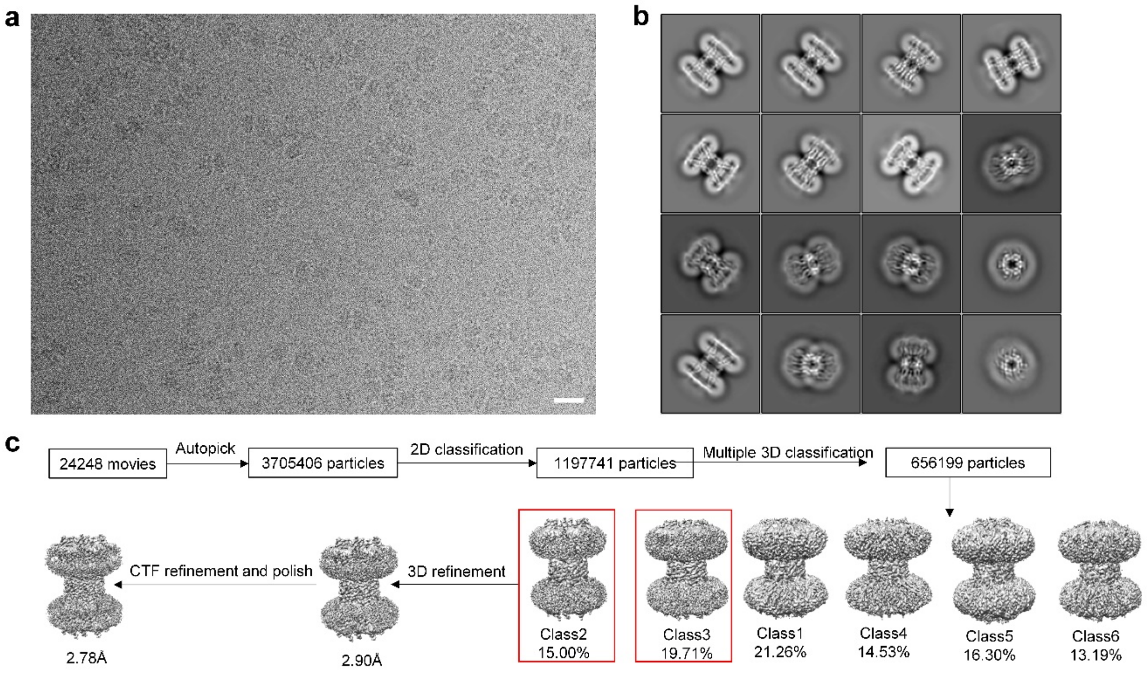

**Extended Data Fig. 4. Cryo-EM data processing pipeline of Cx36-quin in GDN. a,** A representative micrograph of Cx36-quin sample; the scale bar corresponds to 20 nm. **b,** representative 2D classes. **c,** Cryo-EM data process scheme for 3D reconstitution of Cx36-quin in GDN.

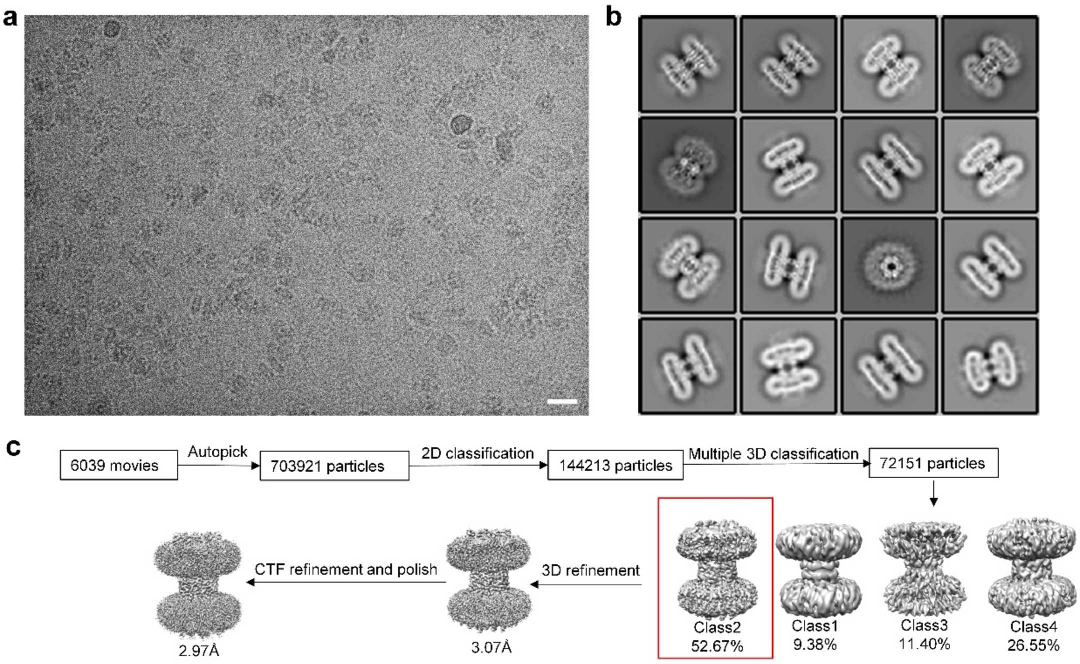

**Extended Data Fig. 5. Cryo-EM data processing pipeline of Cx36-quid in GDN. a,** A representative micrograph of Cx36-quid sample; the scale bar corresponds to 20 nm. **b,** representative 2D classes. **c,** Cryo-EM data process scheme for 3D reconstitution of Cx36-quid in GDN.

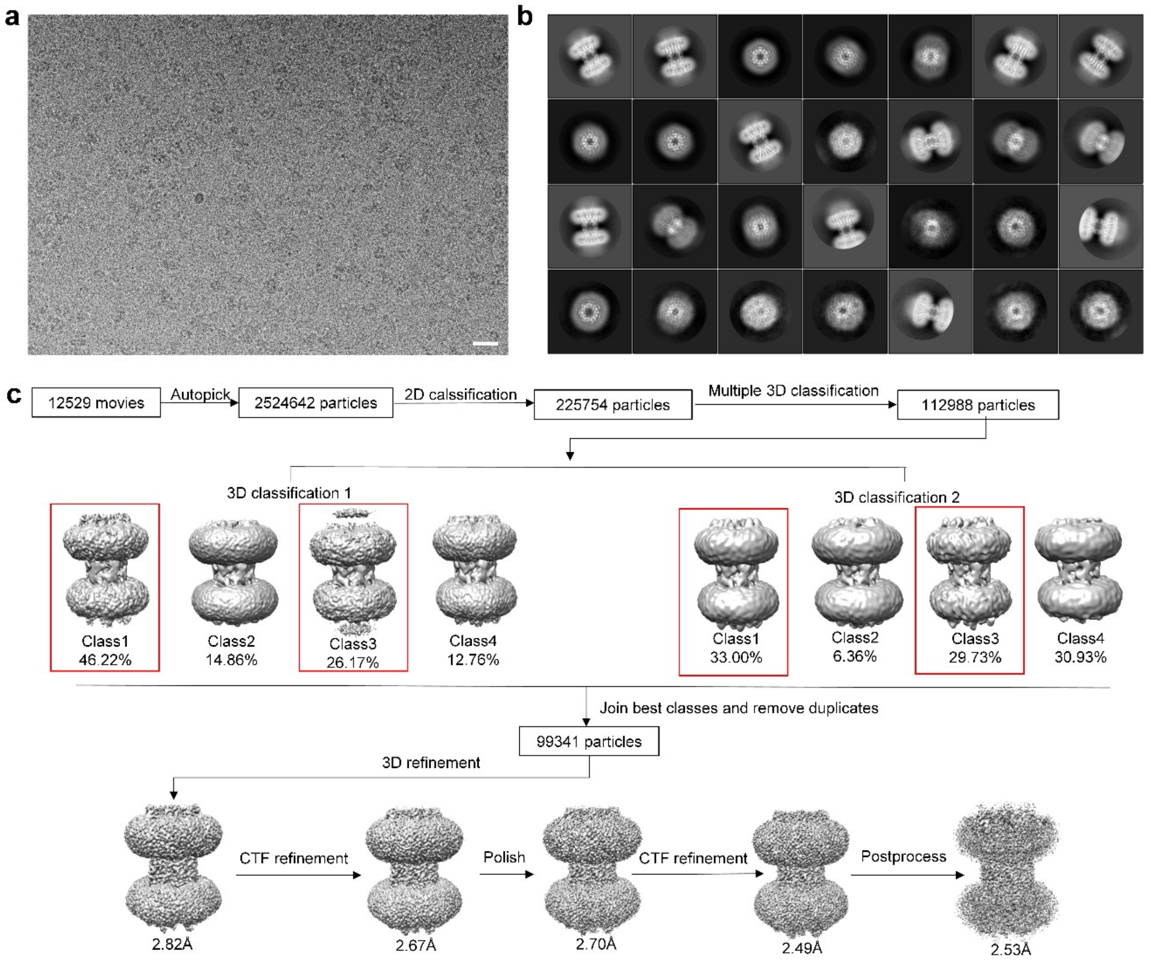

**Extended Data Fig. 6. Cryo-EM data processing pipeline of apo-Cx36 in GDN. a,** A representative micrograph of apo-Cx36 sample; the scale bar corresponds to 20 nm. **b,** representative 2D classes. **c,** Cryo-EM data process scheme for 3D reconstitution of Cx36 in GDN.

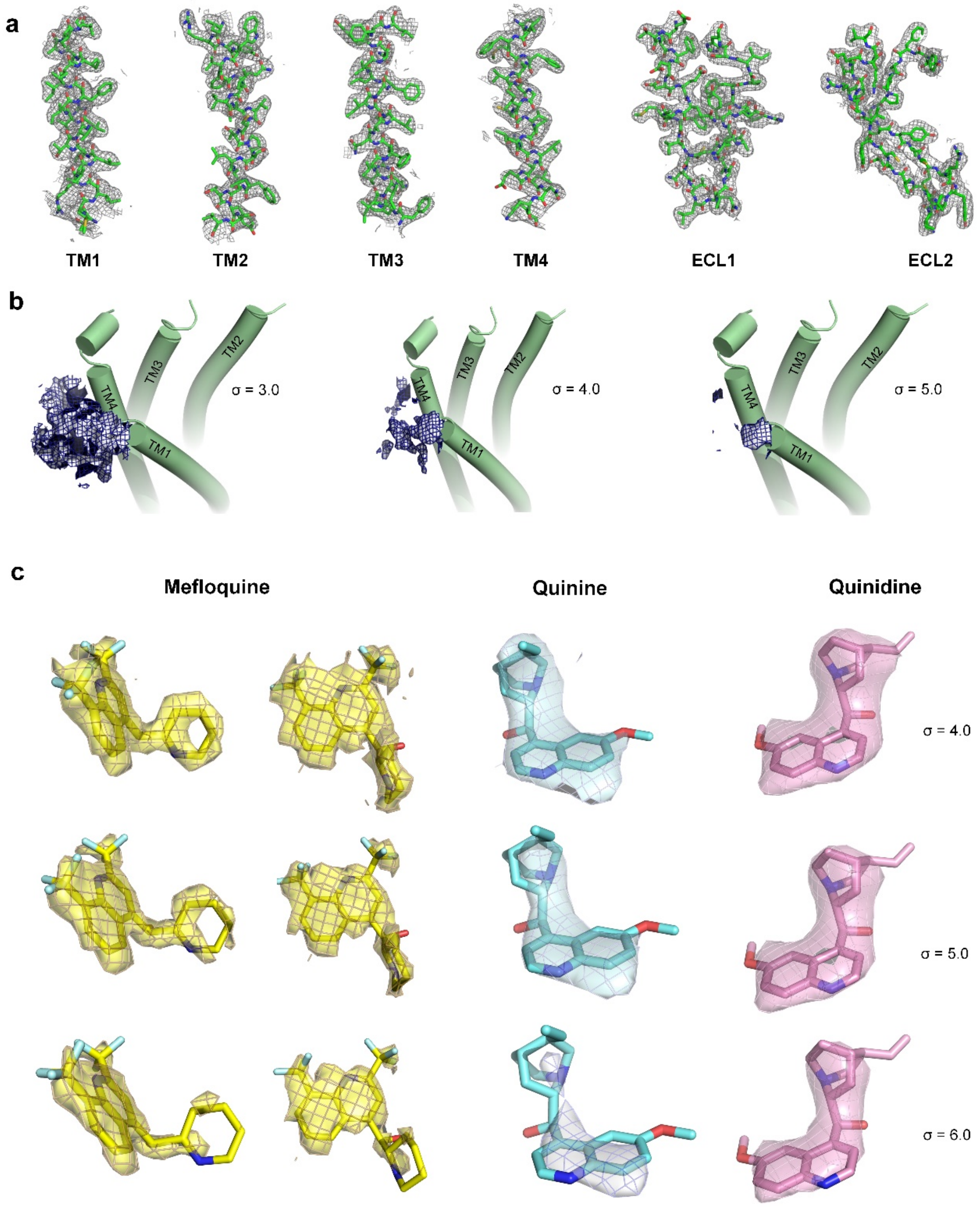

**Extended Data Fig. 7. Cryo-EM density map features of Cx36 and drugs. a,** Representative cryo-EM density map features of Cx36 gap junction channel (GJC) with mefloquine, including the transmembrane (TM) helixes and extracellular loop 1 (ECL1) and extracellular loop 2 (ECL2). **b,** Cryo-EM density map features of N-terminal at σ levels 3.0, 4.0, 5.0. **c,** Cryo-EM density map features of mefloquine (viewed at two angles, illustrating the well-resolved densities corresponding to the rings), quinine and quinidine at σ levels 4.0, 5.0, 6.0 each.

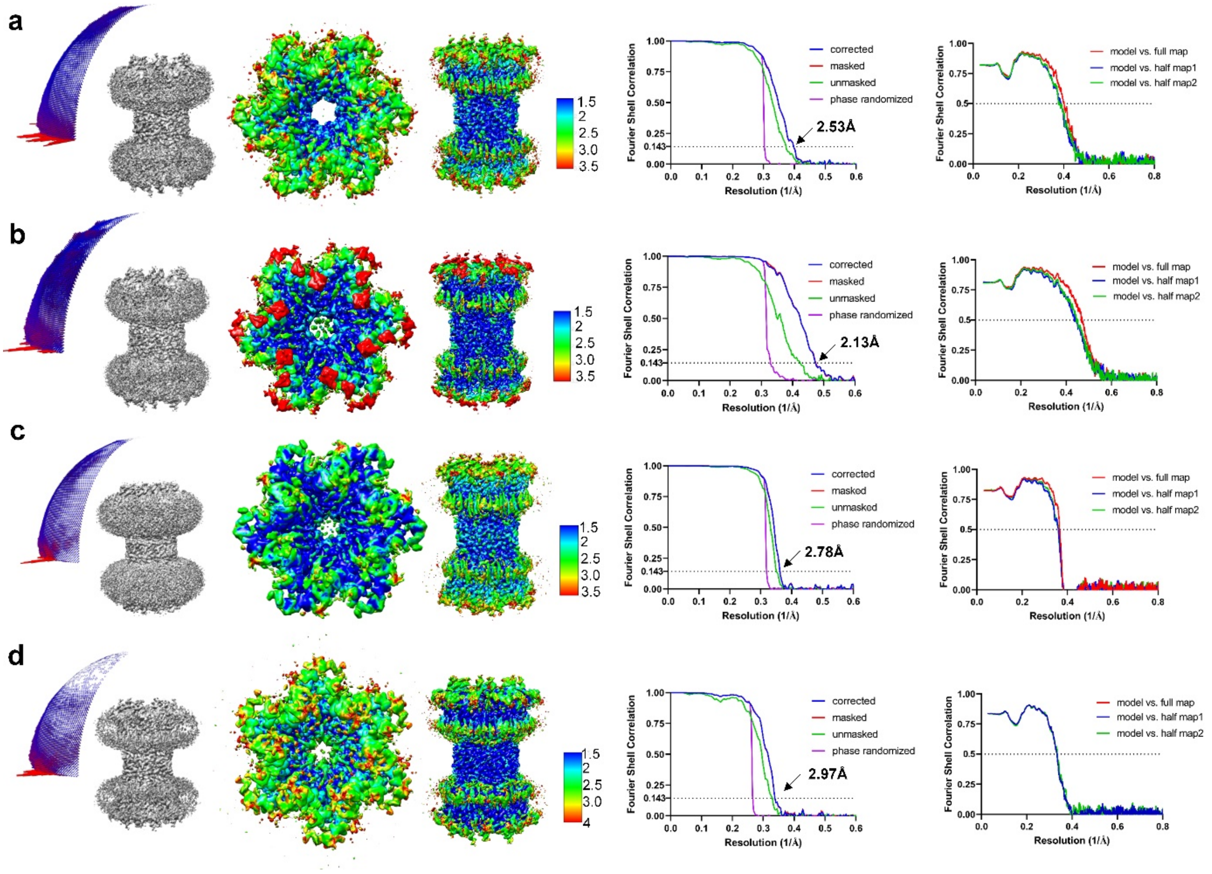

**Extended Data Fig. 8. Angular distribution, local resolution maps and Fourier shell correlation (FSC) plots of Cx36 with and without drugs bound. a,** apo-Cx36. **b,** Cx36-mfq. **c,** Cx36-quin. **d,** Cx36-quid.

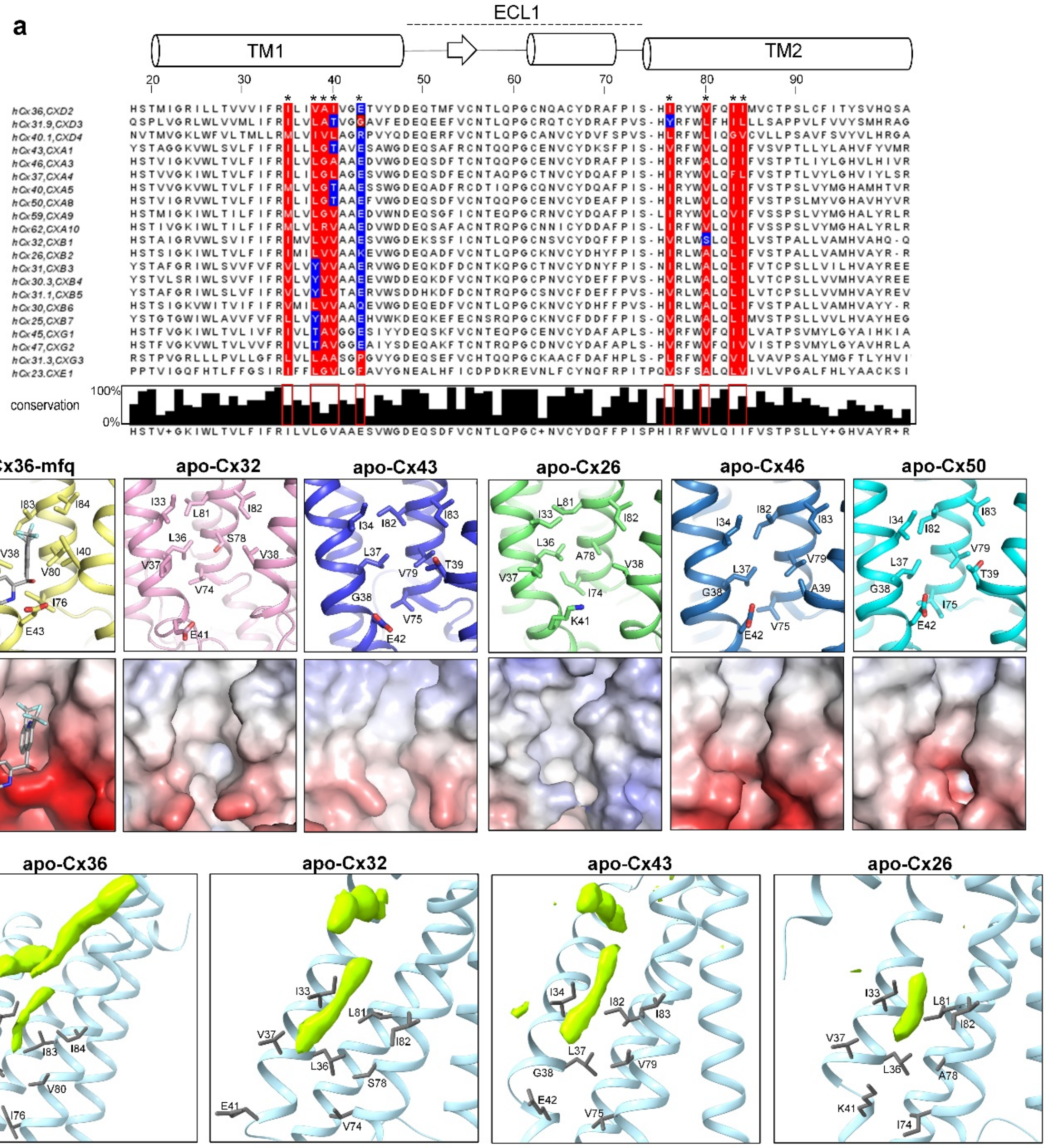

**Extended Data Fig. 9. Amino acid sequence alignment and structural comparison in connexin families. a,** Amino acid sequence alignment of all human connexins in the mefloquine binding site. Sequences are aligned with CLUSTALW [27]. Secondary structures and numbering of residues of Cx36 are represented at the top. The hydrophobic and hydrophilic residues are shaded in red and blue, respectively. Asterisks indicate ligand-binding residues in Cx36. The figure is created with jalview [28] and manually modified. **b,** structural comparison of connexins in the mefloquine binding site. Mefloquine-bound structures of Cx36, Cx32 (PDB 7ZXM), Cx43 (PDB 7Z22) and nonligand-bound structures of Cx26 (PDB 2ZW3), Cx46 (PDB 7JKC), Cx50 (PDB 7JJP) are used. Each panel below shows the electrostatic surface potential representation of binding pocket. **c,** Densities of lipids shown in green are present in the ligand binding sites of Cx36, Cx32 (EMD 15010), Cx43 (EMD 14455), Cx26 (EMD 13938).

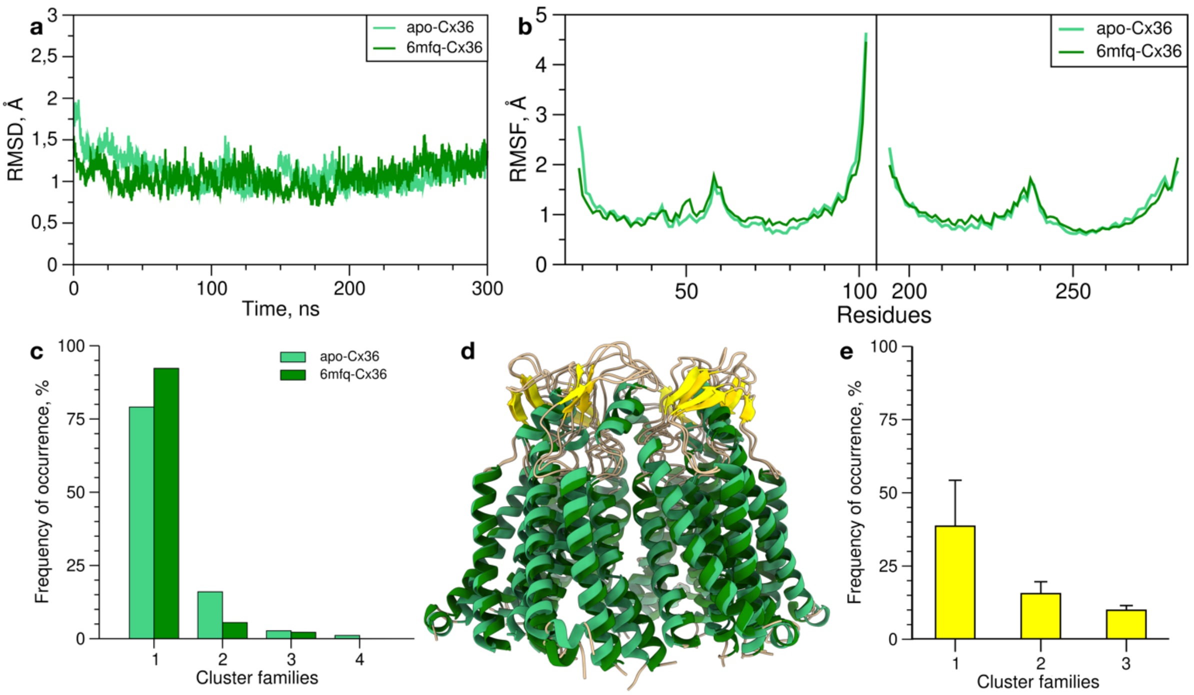

**Extended Data Fig. 10. MD simulations of the *apo-Cx36* and *6mfq-Cx36* systems. a,** A plot reporting the average RMSD of the secondary structure Cα atoms of Cx36 hexamer; **b,** A plot reporting the average per-residue RMSF of each Cx36 monomer. The left panel covers the residues from S19 to A102, whereas the right panel covers the residues from G194 to L282; **c,** Histograms displaying the frequency of occurrence of each cluster family in the *apo-Cx36* and *6mfq-Cx36* MD simulations; **d,** Superimposition of the centroids of the most relevant cluster families measured for the *apo-Cx36* and *6mfq-Cx36* simulations. For *apo-Cx36*, α-helices are displayed as green ribbons and the ß-helices as yellow arrows. For *6mfq-Cx36*, α-helices are displayed as dark green ribbons and the ß-helices as gold arrows; **e,** Histogram displaying the frequency of occurrence of mefloquine's cluster families in the *6mfq-Cx36* MD simulation.

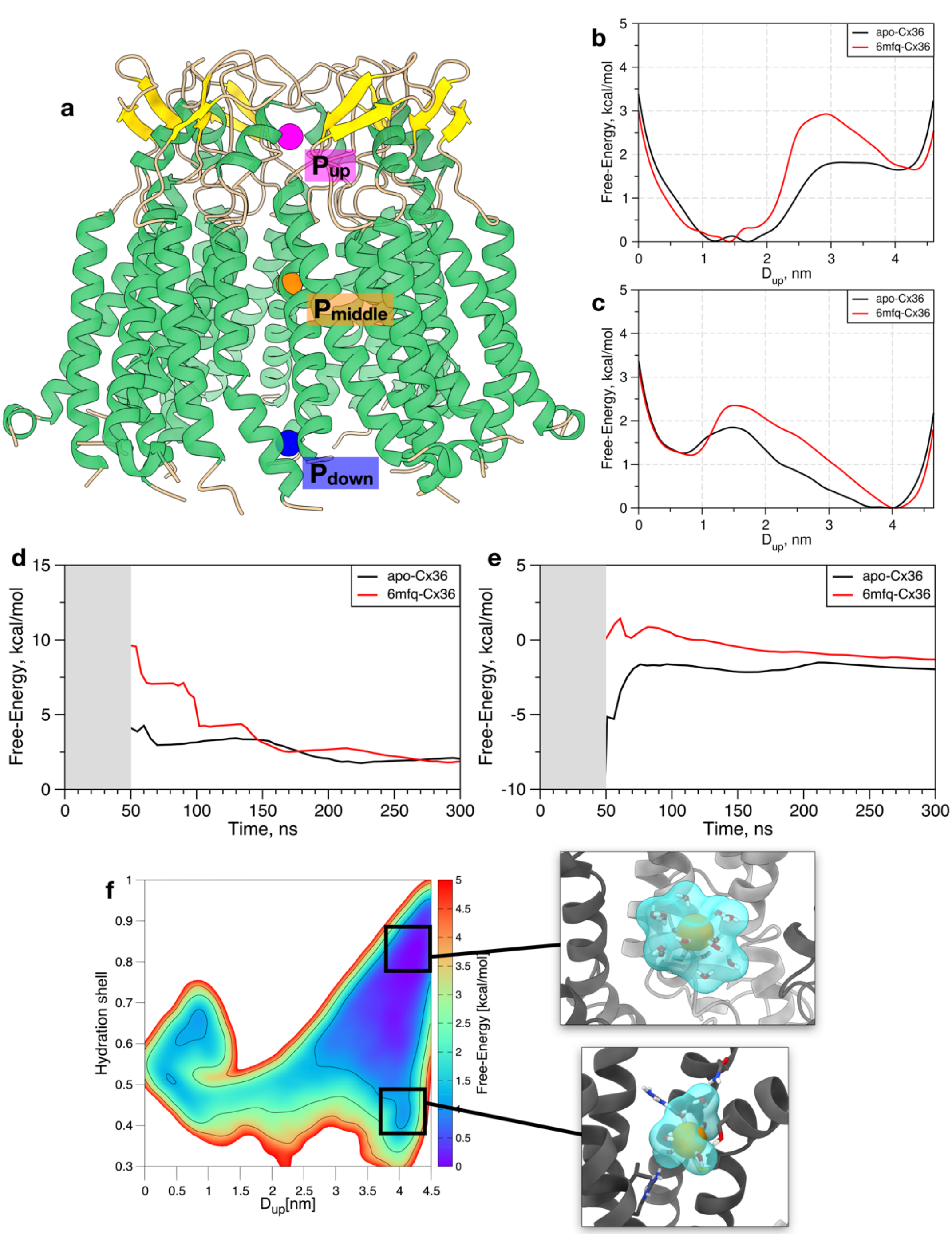

**Extended Data Fig. 11. Details of the OPES simulations carried out on the *apo-Cx36* and *6mfq-Cx36* systems. a,** A schematic depiction of a set of dummy atoms employed to carry out the OPES simulations. The dummy atoms $P_{up}$, $P_{middle}$, and $P_{down}$ are represented as spheres and colored in magenta, orange, and blue, respectively. The α-helices and ß-helices of Cx36 are colored in green and yellow, respectively. **b-c,** Free-energy profiles of the translation of $K^+$ (**b**) and $Cl^-$ (**c**) along Cx36's pore in the *apo-Cx36* and *6mfq-Cx36* OPES simulations. **d-e,** ΔG of translation as function of the simulation time for the $K^+$ (**d**) and $Cl^-$ (**e**) ions in the OPES simulations performed on the *apo-Cx36* and *6mfq-Cx36* systems; **f,** Free-energy surfaces associated with Cl- permeation across the Cx36 hexamer in the *apo-Cx36*. The insets show two representative frames of the different hydration state of Cl- ion in the intracellular portion of the Cx36 hexamer.

**Extended Data Table 1.** Cryo-EM analysis and statistics

| | | | | |
|---|---|---|---|---|
| **Data collection** | | | | |
| Sample | apo-Cx36 | Cx36-mfq | Cx36-quin | Cx36-quid |
| Instrument | FEI Titan Krios / Gatan K3 Summit | | | |
| Voltage (kV) | 300 | | | |
| Electron Dose (e-/Å2) | 50 | 55 | 50 | 55 |
| Defocus range (µm) | -0.5 to -2.5 | | | |
| Pixel size (Å) | 0.65 | 0.66 | 0.66 | 0.66 |
| Number of particles | 99341 | 102585 | 231305 | 39915 |
| FSC threshold 0.143 | 2.53 | 2.13 | 2.78 | 2.97 |
| **Refinement** | | | | |
| Model resolution FSC threshold 0.5 | 2.49 | 2.14 | 2.73 | 2.90 |
| Map sharpening B-factor (Å) | -84.6697 | -51.3034 | -103.162 | -90.2861 |
| Map CC | 0.96 | 0.94 | 0.81 | 0.84 |
| **Model composition** | | | | |
| Protein residues/ligands/water | 2076/0/90 | 2076/12/162 | 2076/12/0 | 2076/12/0 |
| Bond length r.m.s.d (Å) | 0.003 | 0.002 | 0.004 | 0.003 |
| Bond angle r.m.s.d (°) | 0.461 | 0.508 | 0.574 | 0.508 |
| **Validation** | | | | |
| MolProbity score | 1.07 | 1.17 | 1.31 | 1.27 |
| Clash score | 2.82 | 2.43 | 3.68 | 3.57 |
| Rotamer outlier (%) | 0.53 | 1.55 | 1.60 | 1.44 |
| Ramachandran plot | | | | |
| Favoured (%) | 98.62 | 99.31 | 99.41 | 99.31 |
| Allowed (%) | 1.38 | 0.69 | 0.59 | 0.69 |
| Disallowed (%) | 0.00 | 0.00 | 0.00 | 0.00 |

**Extended Data Table 2.** Crossing events of the ions $K^+$ and $Cl^-$ in the MD simulations carried out on the *apo-Cx36* and *6mfq-Cx36* systems.

| | **apo-Cx36** | **6mfq-Cx36** |
|---|---|---|
| **$K^+$ ions** | 30 | 7 |
| **$Cl^-$ ions** | 6 | 2 |
